# Evaluation of GRCh38 and de novo haploid genome assemblies demonstrates the enduring quality of the reference assembly

**DOI:** 10.1101/072116

**Authors:** Valerie A. Schneider, Tina Graves-Lindsay, Kerstin Howe, Nathan Bouk, Hsiu-Chuan Chen, Paul A. Kitts, Terence D. Murphy, Kim D. Pruitt, Françoise Thibaud-Nissen, Derek Albracht, Robert S. Fulton, Milinn Kremitzki, Vince Magrini, Chris Markovic, Sean McGrath, Karyn Meltz Steinberg, Kate Auger, Will Chow, Joanna Collins, Glenn Harden, Tim Hubbard, Sarah Pelan, Jared T. Simpson, Glen Threadgold, James Torrance, Jonathan Wood, Laura Clarke, Sergey Koren, Matthew Boitano, Heng Li, Chen-Shan Chin, Adam M. Phillippy, Richard Durbin, Richard K. Wilson, Paul Flicek, Deanna M. Church

## Abstract

The human reference genome assembly plays a central role in nearly all aspects of today’s basic and clinical research. GRCh38 is the first coordinate-changing assembly update since 2009 and reflects the resolution of roughly 1000 issues and encompasses modifications ranging from thousands of single base changes to megabase-scale path reorganizations, gap closures and localization of previously orphaned sequences. We developed a new approach to sequence generation for targeted base updates and used data from new genome mapping technologies and single haplotype resources to identify and resolve larger assembly issues. For the first time, the reference assembly contains sequence-based representations for the centromeres. We also expanded the number of alternate loci to create a reference that provides a more robust representation of human population variation. We demonstrate that the updates render the reference an improved annotation substrate, alter read alignments in unchanged regions and impact variant interpretation at clinically relevant loci. We additionally evaluated a collection of new de novo long-read haploid assemblies and conclude that while the new assemblies compare favorably to the reference with respect to continuity, error rate, and gene completeness, the reference still provides the best representation for complex genomic regions and coding sequences. We assert that the collected updates in GRCh38 make the newer assembly a more robust substrate for comprehensive analyses that will promote our understanding of human biology and advance our efforts to improve health.

## Introduction

The human reference genome assembly remains a critical resource for the biological and clinical research communities (International Human Genome Sequencing Consortium 2004; Lander et al. 2001). It is distinguished from the growing number of human genome assemblies in public databases by virtue of its long contig and scaffold N50s, high base-pair accuracy, and robust representations of repetitive and segmentally duplicated genomic regions, all of which reflect the clone-based assembly approach and Sanger sequencing methods that were the basis of its generation. Assembled from the DNA of multiple donors, it was intended to provide representation for the pan-human genome, rather than a single individual or population group and is a mosaic of haplotypes whose boundaries coincide with the underlying clone boundaries.

A revision to the assembly model, first used in the prior version of the reference, GRCh37 (GCA_000001405.1), expanded the ability of the reference assembly to represent the extent of structural variation and population genomic diversity whose discovery it facilitated (Church et al. 2011; 1000 Genomes Project Consortium et al. 2015; International HapMap Consortium 2005; Sudmant et al. 2010; Kidd et al. 2008). The introduction of alternate loci scaffolds enabled GRCh37 to include additional sequence representations for the highly variant MHC region, as well as the divergent haplotypes of the MAPT and UGT2B loci, while retaining the linear chromosome representations familiar and intuitive to most users (Zody et al. 2008; Xue et al. 2008; Horton et al. 2008). A second feature of the updated model, assembly patches, permitted subsequent corrections and addition of new sequence representations to the GRCh37 assembly without changing the chromosome sequences or coordinates upon which an increasing volume of data were being mapped (1000 Genomes Project Consortium et al. 2015; Pierson et al. 2015; Zook et al. 2014). The assembly model remains for GRCh38, the current reference version. Together, these features of the assembly model helped ensure that the human reference assembly would continue to present the most accurate representation of the human genome possible while providing a stable substrate for large-scale analysis.

The GRCh37 assembly underwent thirteen patch releases in the period from 2009 to 2013 (GCA_000001405.2-GCA_000001405.14). Despite the availability of these sequences in public databases, their use has been limited by the inability of common bioinformatics file formats and tool chains to manage the allelic duplication they introduce, as well as by their constrained representation in popular genome browsers (Church et al. 2015). In addition, the patches represented only a subset of the assembly updates made by the Genome Reference Consortium (GRC). Thus, coordinate changing assembly updates remain essential for users to access the full suite of assembly improvements, despite the challenge of transporting data and results to the new assembly (Zhao et al. 2014; Hickey et al. 2013).

In producing GRCh38, we of the Genome Reference Consortium (GRC) placed special emphasis on addressing the following types of assembly issues found in GRCh37: (1) resolution of tiling path errors and gaps associated with complex haplotypes and segmental duplications; (2) base-pair level updates for sequencing errors; (3) addition of “missing” sequences, with an emphasis on paralogous sequences and population variation; and (4) providing sequence representation for genomic features, such as centromeres and telomeres. Making these updates involved the use of bioinformatics and experimental resources and techniques not previously available. We will demonstrate how the new approaches used in this effort result in a human reference genome assembly that is more contiguous and complete than ever before, and which provides better gene and variant representation than GRCh37, features critical to both basic research and clinical uses of the assembly. We will also show how assembly updates in GRCh38 impact analyses throughout the genome, even in regions that are unchanged between the two assemblies. Together, these analyses suggest adoption of the new assembly will have a positive impact on both genome-wide analysis as well as regional analysis.

With long range sequencing and assembly technologies making the generation of highly contiguous whole genome de novo assemblies possible, the overall value of GRCh38 and the human reference genome assembly in general, must now also be considered (Chaisson et al. 2015b). The reference assembly is not just a substrate for alignment, but is also the coordinate system upon which we annotate our biological knowledge. Several recently published individual human de novo assemblies have been favorably compared to GRCh38 with respect to continuity metrics, and while they each contain sequence not present in the reference assembly, none yet surpass the global quality of GRCh38 (Cao et al. 2015; Li et al. 2010; Pendleton et al. 2015; Steinberg et al. 2014; Berlin et al. 2015). Such assemblies are often suggested as sequence sources for use in closure of reference assembly gaps, while other studies have called for one or more individual genomes to replace the reference (Rosenfeld et al. 2012). To address these issues, we used a range of assembly metrics to generate and evaluate a collection of de novo assemblies derived from the same sequence data and assembled using different algorithms and/or parameters, with respect to each other and in comparison to GRCh38. The de novo assemblies were derived from the essentially haploid complete hydatidiform mole samples CHM1 and CHM13 (Steinberg et al. 2014; Fan et al. 2002). The absence of allelic variation in these samples simplifies their assembly relative to diploid samples, as only repeat content remains to confound assembly construction. Consistent with prior multi-assembly evaluations, these analyses showed that no single assembly parameter or algorithm creates a best whole genome assembly across all metrics evaluated (Vezzi et al. 2012; Bradnam et al. 2013; Earl et al. 2011). More notably, they revealed that while de novo assemblies can now approach reference quality for some metrics, for others they still lag considerably behind GRCh38. In doing so, they further reveal that the use of sequence from such assemblies to close gaps or add to the reference assembly may not always be desirable. Not only do our data demonstrate a continued role and relevance for the current human genome reference assembly, they emphasize the need for continued development in the fields of sequencing and assembly so that the human reference genome of the future exhibits the necessary all-around quality essential to fulfill its many roles in an ever-expanding set of analyses.

## Results

### Assembly Updates

Upon the release of GRCh37.p13 in June 2013, the cumulative set of 204 patch scaffolds covered 3.15% of the chromosome assemblies and included more than 7 Mb of novel sequence, and met previously defined GRC criteria for the trigger of a major assembly release (Church et al. 2011). We submitted GRCh38, a coordinate changing update of the human reference assembly, to the International Nucleotide Sequence Database Collaboration (INSDC) in December, 2013 (GCA_000001405.15). As the reference remains under active curation, we have subsequently provided quarterly GRCh38 patch releases, which do not affect the chromosome coordinates, the latest of which was GRCh38.p8 (GCA_000001405.23). The initial GRCh38 release represents the resolution of more than 1000 issues reported to the GRC tracking system, spanning all chromosomes and encompassing a variety of problem types, including gaps, component and tiling path errors and variant representation (Figure 1 and (https://www.ncbi.nlm.nih.gov/projects/genome/assembly/grc/human/issues/). As in prior assembly updates, we sought to use finished, clone-based components for assembly updates wherever possible, due to their high per-base accuracy and haploid representation of actual human sequence. With >95% of the chromosome total sequence, and 98% of non-centromeric sequence, derived from genomic clone components, the GRCh38 reference assembly chromosomes continue to provide a mosaic haploid representation of the human genome, rather than a consensus haploid representation. The sequence contribution from RP11, an anonymous male donor of likely African-European admixed ancestry, remains dominant (~70%), but has decreased by ~1.5% relative to the prior assembly version (Figure 1) (Green et al. 2010).

**Figure 1.**
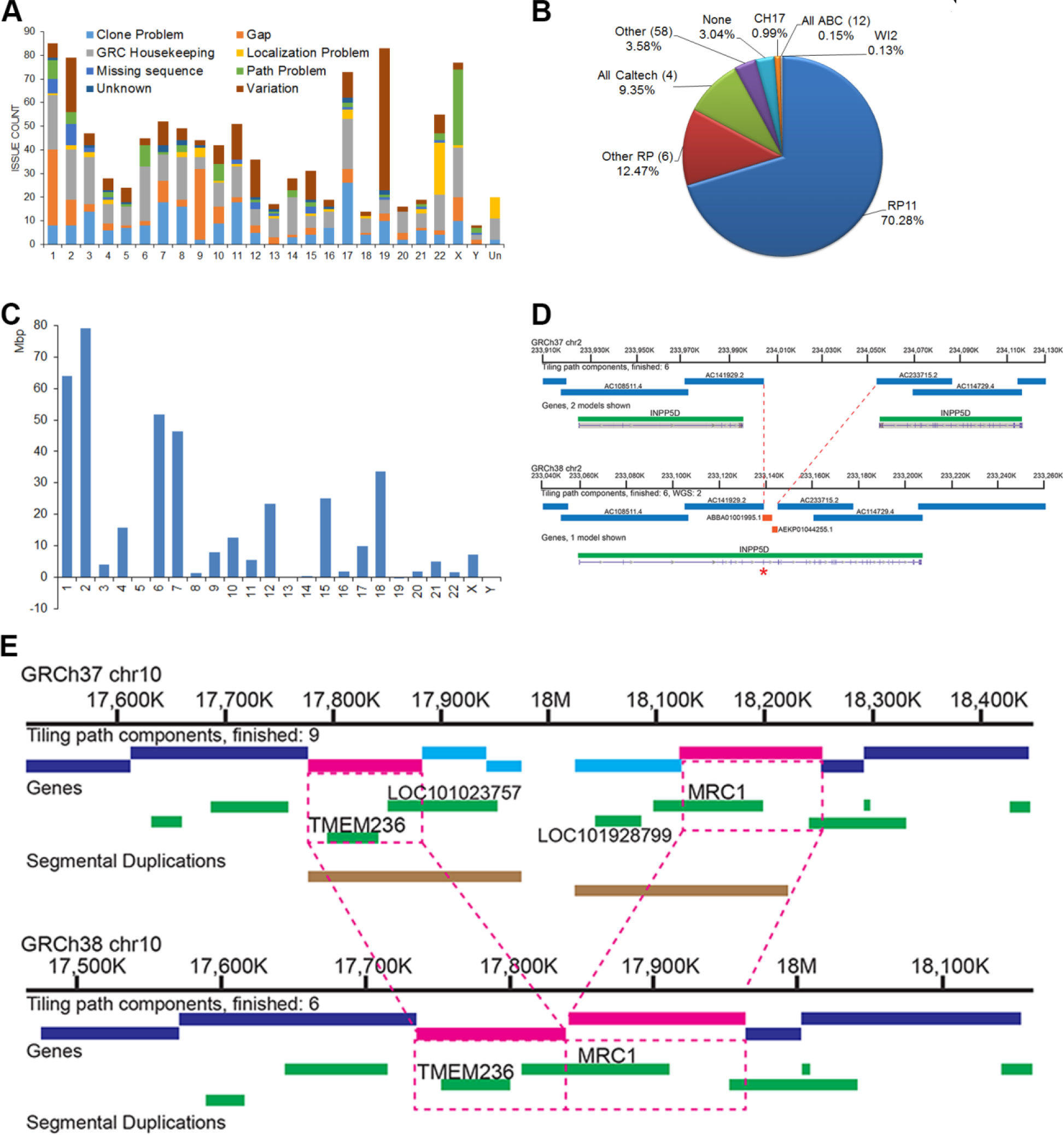
Summary of GRCh38 updates. (A) Chart showing issues resolved for GRCh38 on each chromosome, by issue type. (B) Genomic library composition of GRCh38 primary assembly unit sequence. In some cases, genomic clone libraries have been grouped by creator (RP, Caltech, ABC). The count of samples contributing to groups is shown. None = WGS or PCR sequence. Clone libraries are described in the NCBI Clone DB (https://www.ncbi.nlm.nih.gov/clone/) (C) Changes in placed scaffold N50 length from GRCh37 to GRCh38. Changes on chrs 5, 13, 19 and Y are < 55 Kbp each. (D) Addition of WGS components (orange bars) resolves a GRCh37 gap, consolidating the split annotation of *INPP5D* and restoring a missing exon (red asterisk) in GRCh38. The default 50 Kbp gap in GRCh37 greatly overestimates the actual amount of missing sequence (~ 6 Kbp). (E) Schematic of a curated collapse in GRCh38 chr 10. Clones from two incompatible haplotypes (pink and light blue) were mixed in the GRCh37 tiling path, creating a false gap and segmental duplication involving the single copy genes *TMEM236* and *MRC1* (top). In GRCh38 (bottom), clones from the blue haplotype have been eliminated (~200 Kbp), closing the gap and providing the correct gene content.

Table 1 summarizes the GRCh38 assembly statistics of length, N50 and gaps relative to GRCh37 and several recently generated de novo assemblies. The GRCh38 assembly is longer and more contiguous than prior reference assembly versions (Figure 1, Table 1 and see https://www.ncbi.nlm.nih.gov/projects/genome/assembly/grc/human/data/). Although the total number of reference assembly gaps increased, increases occur when sequence added into a pre-existing gap is not contiguous with either gap edge or when sequence additions are comprised of scaffolded WGS contigs. The increase in gap count in GRCh38 is largely attributable to the replacement of the single centromere gap in each chromosome with scaffolds of modeled sequence (described below), and WGS sequences flank more unspanned gaps and spanned gaps in GRCh38 than in GRCh37 (Supplemental Table S1).

**Table 1.** Comparison of Assembly Statistics

| Assembly Short Name | GenBank Accession | Total Length | Contig N50 | Scaffold N50 | Gap number | Gap Length | QV |
| --- | --- | --- | --- | --- | --- | --- | --- |
| GRCh38* | GCA_000001405.15 | 3,209,286,105 | 56,413,054 | 67,794,873 | 349<sup>^</sup><br>526<sup>\$</sup><br>124<sup>#</sup> | 159,970,007 | ND |
| GRCh37* | GCA_000001405.1 | 3,137,144,693 | 38,508,932 | 46,395,641 | 86<sup>^</sup><br>271<sup>\$</sup><br>100<sup>#</sup> | 239,850,738 | ND |
| CHM1_1.1 | GCA_000306695.2 | 3,037,866,619 | 143,936 | 50,362,920 | 225<sup>^</sup><br>40,665<sup>\$</sup> | 210,229,812 | ND |
| CHM1_CA_P6 | GCA_001307025.1 | 2,939,630,703 | 20,609,304 | NA | 0 | NA | 42.29 |
| CHM1_FC_P6 | GCA_001297185.1 | 2,996,426,293 | 26,899,841 | NA | 0 | NA | 44.64 |
| CHM13_CA1 | GCA_000983465.1 | 3,061,240,732 | 13,331,528 | NA | 0 | NA | 41.21 |
| CHM13_CA2 | GCA_001015355.1 | 3,028,917,871 | 19,357,701 | NA | 0 | NA | 39.86 |
| CHM13_CA3 | GCA_000983475.1 | 2,996,416,935 | 5,550,336 | NA | 0 | NA | 42.89 |
| CHM13_CA4 | GCA_001015385.3 | 3,065,003,163 | 12,252,446 | NA | 0 | NA | 41.27 |
| CHM13_FC | GCA_000983455.2 | 2,941,135,618 | 10,549,591 | NA | 0 | NA | 43.00 |

\* Values include alternate loci unless noted <sup>^</sup> Scaffold breaking gap <sup>\$</sup> Non-breaking gap (excludes alt loci) <sup>#</sup> Non-breaking gap (alt loci)

The suite of updates provided in the GRCh38 assembly had a positive impact on assembly annotation. Comparison of the NCBI *Homo sapiens* annotation release 105 of GRCh37.p13 (https://www.ncbi.nlm.nih.gov/genome/annotation_euk/Homo_sapiens/105/) and annotation release 106 of GRCh38 (https://www.ncbi.nlm.nih.gov/genome/annotation_euk/Homo_sapiens/106/) shows an increase in the numbers of genes and protein coding transcripts, with a concomitant decrease in partially represented coding sequences and transcripts split over assembly gaps (Table 2). Because the transcript content of these two annotation releases was not identical, we also aligned two large public annotation sets (GENCODE23 (basic) and RefSeq71) to the GRCh37 and GRCh38 full assemblies to gauge the impact of improvements on gene representation (O’Leary et al. 2016; Harrow et al. 2012). Similar to the previously described comparison, in GRCh38 we find that both annotation sets show increases in overall transcript alignments with a substantial decrease in split and low quality transcript alignments (Table 3, Supplemental Worksheet S1). We looked at the intersection of the transcripts with problematic alignments with two clinically relevant genes lists: a set of genes enriched for de novo loss of function mutations identified in Autism Spectrum Disorder (n=1003) (Samocha et al. 2014) and a collection of genes preliminarily proposed for the development of a medical exome kit (n=4623) (https://www.genomeweb.com/sequencing/emory-chop-harvard-develop-medical-exome-kit-complete-coverage-5k-disease-associ). Among the set of RefSeq transcripts with problematic alignments to GRCh37, we observed 6 gene overlaps with the former and 14 with the latter, while we found 6 and 22 for the GENCODE cohort (Supplemental Worksheet S1). The majority of these genes are no longer associated with transcript alignment issues in GRCh38, suggesting the newer assembly is a better substrate for clinical studies.

**Table 2.** Summary of RefSeq Annotation Releases 105 and 106

| Feature | NCBI Annotation Release 105* |  |  | NCBI Annotation Release 106^ |  |  |
| --- | --- | --- | --- | --- | --- | --- |
|  | GRCh37.p13 |  |  | GRCh38 |  |  |
|  | Full Assembly | Primary Assembly | All Alt Loci | Full Assembly | Primary Assembly | All Alt Loci |
| **Genes and pseudogenes** | 40,158 | 39,947 | 428 | 41,722 | 41,566 | 1,981 |
| **mRNAs** | 67,517 | 64,734 | 1,360 | 69,826 | 67,793 | 3,408 |
| **Other RNAs** | 15,063 | 14,151 | 443 | 17,857 | 16,914 | 1,152 |
| **CDSs** | 68,035 | 65,099 | 1,360 | 70,368 | 68,177 | 3,564 |

\* Entrez query date: Aug 3 2013 (42,339 known RefSeqs (NM\_/NR\_) https://www.ncbi.nlm.nih.gov/genome/annotation_euk/Homo_sapiens/105/
^ Entrez query date: Jan 17 2014 (45,911 known RefSeqs (NM\_/NR\_) https://www.ncbi.nlm.nih.gov/genome/annotation_euk/Homo_sapiens/106/

**Table 3.**
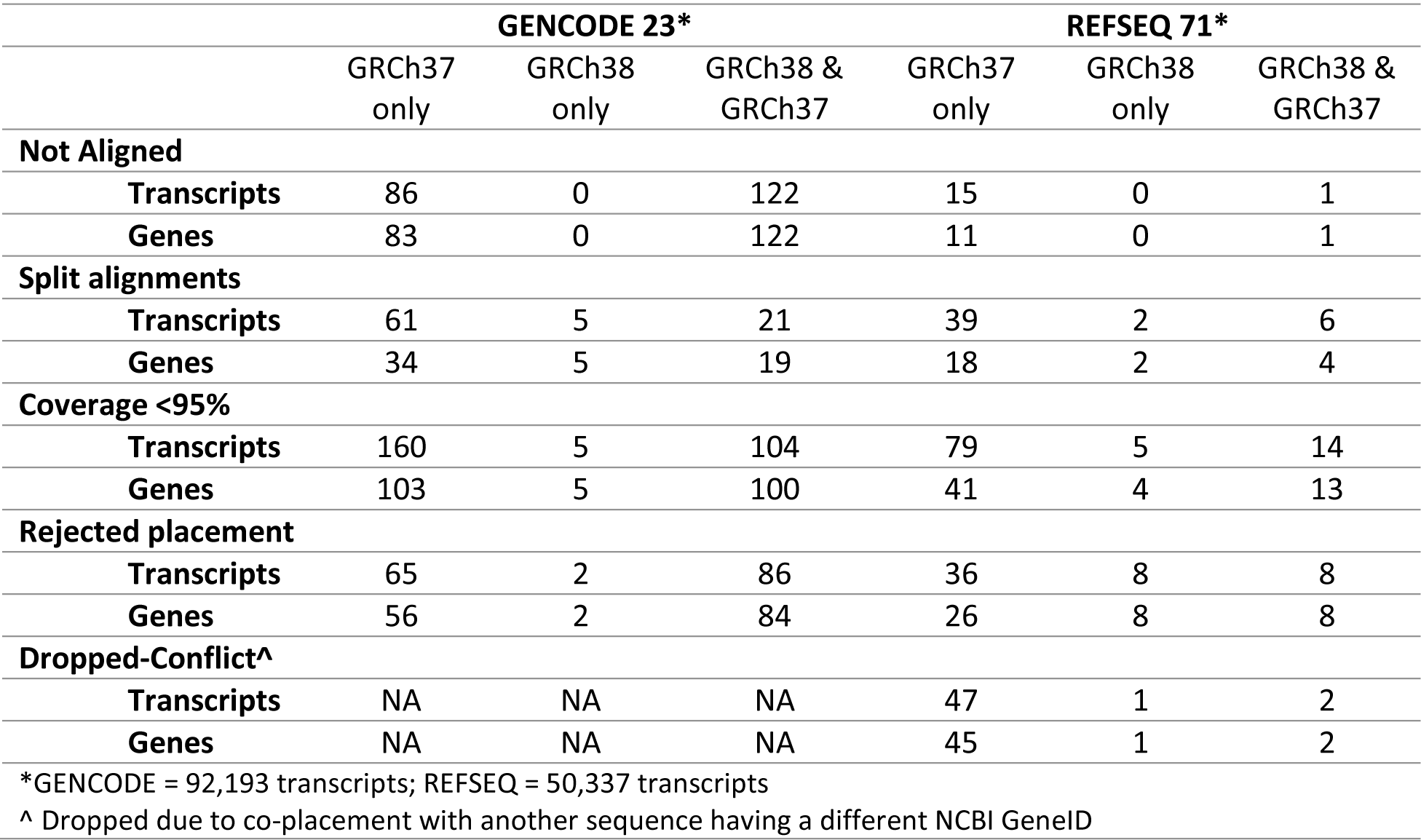
GENCODE 23 and REFSEQ 71 Alignments to GRCh37 and GRCh38

|  | GENCODE 23* |  |  | REFSEQ 71* |  |  |
| --- | --- | --- | --- | --- | --- | --- |
|  | GRCh37 only | GRCh38 only | GRCh38 & GRCh37 | GRCh37 only | GRCh38 only | GRCh38 & GRCh37 |
| **Not Aligned** |  |  |  |  |  |  |
| **Transcripts** | 86 | 0 | 122 | 15 | 0 | 1 |
| **Genes** | 83 | 0 | 122 | 11 | 0 | 1 |
| **Split alignments** |  |  |  |  |  |  |
| **Transcripts** | 61 | 5 | 21 | 39 | 2 | 6 |
| **Genes** | 34 | 5 | 19 | 18 | 2 | 4 |
| **Coverage <95%** |  |  |  |  |  |  |
| **Transcripts** | 160 | 5 | 104 | 79 | 5 | 14 |
| **Genes** | 103 | 5 | 100 | 41 | 4 | 13 |
| **Rejected placement** |  |  |  |  |  |  |
| **Transcripts** | 65 | 2 | 86 | 36 | 8 | 8 |
| **Genes** | 56 | 2 | 84 | 26 | 8 | 8 |
| **Dropped-Conflict<sup>^</sup>** |  |  |  |  |  |  |
| **Transcripts** | NA | NA | NA | 47 | 1 | 2 |
| **Genes** | NA | NA | NA | 45 | 1 | 2 |

\*GENCODE = 92,193 transcripts; REFSEQ = 50,337 transcripts <sup>^</sup> Dropped due to co-placement with another sequence having a different NCBI GeneID

### Centromeres

A major change in the content of the reference genome assembly is the replacement of the 3 Mbp centromeric gaps on all GRCh37 chromosomes with modeled centromeres from the LinearCen1.1 (normalized) assembly, derived from a database of centromeric sequences from the HuRef genome (GCA_000442335.2) (Miga et al. 2014; Levy et al. 2007) (Supplemental Methods). We added the modeled centromeres to the reference assembly to serve as catalysts for analyses of these biologically important and highly variant genomic regions, as annotation targets, and to act as read sinks for centromere-containing reads in mapping analyses (Miga et al. 2015). Consistent with our intent, 21.7% (by length) of the “decoy” sequence used in the 1000 Genomes project to reduce spurious read mapping was identified by RepeatMasker as alpha-satellite centromeric repeat (1000 Genomes Project Consortium et al. 2015) (ftp://ftp.1000genomes.ebi.ac.uk/vol1/ftp/technical/reference/phase2_reference_assembly_sequence/). Each centromere model represents the variants and monomer ordering of the chromosome-specific alpha-satellite repeats in a manner proportional to that observed in the initial read database, but the long-range ordering of repeats is inferred. In contrast to the remainder of the chromosome sequence, in which each underlying clone component represents the actual haplotype of its source DNA, the modeled sequence is not an actual haplotype, but an averaged representation. The GRCh38 modeled centromeres also contain largely unordered and unoriented islands of euchromatic sequences that are taken from the same collection of HuRef sequences, as well as from genomic clones. One such island, in the modeled centromere for chromosome 3, provides reference representation for a *PRIM2* paralog (NCBI gene LOC101930420) that was missing in GRCh37 (Genovese et al. 2013a, 2013b). Due to the modeled nature of these sequence representations, we suggest that variant and other analyses within these regions be treated independently of similar analyses made elsewhere in the genome. We anticipate that these modeled sequences will be updated in future assembly versions as new sequencing and assembly technologies make it possible to provide longer-range representations for these regions.

### Retiling

While a subset of missing sequences are associated with gaps deemed recalcitrant to cloning, segmental duplications or other complex genomic architectures are implicated in most remaining gaps or mis-assemblies (Chaisson et al. 2015a; Sharp et al. 2005; Bailey et al. 2001). In collaboration with various external groups, we identified and investigated reported path issues and associated assembly gaps using a combination of techniques, including optical maps, Strand-Seq, admixture mapping and re-evaluation of component sequences and overlaps (Mueller et al. 2013; Falconer et al. 2012; Teague et al. 2010; Howe and Wood 2015; Genovese et al. 2013a). These analyses uncovered some substantial mis-assemblies in GRCh37 that spanned several megabases and many genes, including the regions at 1q21, 10q11 and a pericentromeric inversion of chromosome 9. While we were able to improve or resolve some path problems through reordering of existing assembly components to match optical maps, we found that other approaches were needed at more complex regions where allelic and paralogous variation made it impossible to confidently define paths with clones representing a mosaic of diploid DNA sources. In these instances, we replaced GRCh37 components with new tiling paths comprised of BAC clones representing the single haplotype of the essentially haploid CHM1 genome, or on chromosome X, with the single haplotype represented in RP11 (Steinberg et al. 2014; Dennis et al. 2012; Mueller et al. 2013). We also re-tiled several genomic loci associated with immune responses (IGK, IGH, LRC-KIR and the cytokine cluster on 17q) with CHM1 clones, replacing the contrived mosaic representations in GRCh37 and prior assembly versions to ensure the reference provided an accurate representation of these clinically important regions (Supplemental Worksheet S2) (Watson et al. 2015, 2013). It is important to note that these new representations may not always be common across any or all populations. Wherever possible, we preserved the assembly representation of genes for which the CHM1 haplotype is deleted by adding components containing these genes to alternate loci scaffolds. Resolution of tiling path issues and assembly gaps is not always accompanied by sequence addition or replacement. For example, we removed 3 components on chromosome 10, representing approximately 200 Kbp of falsely duplicated sequence, to close a gap and correct gene representation (Figure 1). Ongoing reference assembly curation efforts include providing haplotype resolved paths at other complex loci, such as the Prader-Willi and flanking regions at 15q11-13 (Antonacci et al. 2014).

### Paralogous sequence additions

In the course of closing gaps and correcting path errors, we focused on providing reference assembly representation for previously missing human-specific and paralogous sequences. More than 100 segmentally duplicated regions have been estimated to be underrepresented in prior versions of the reference assembly (Sudmant et al. 2010). We have previously shown that an incomplete reference assembly can lead to incorrect mapping of reads (Church et al., 2011) which could subsequently lead to misidentifying paralogous sequence variants as allelic sequence variants. With reported regions as a guide, we used whole genome maps, admixture mapping, and FISH and alignment analyses to resolve mis-assemblies and identify and localize components in the assembly. To evaluate our efforts, we analyzed NCBI assembly-assembly alignments of GRCh37 and GRCh38 to determine the relative extents of expansion and collapse in the two assemblies. The NCBI alignment protocol produces outputs that include both reciprocal best hits and non-reciprocal best hits (Steinberg et al. 2014; Kitts et al. 2016). For a given assembly in an alignment pair, genomic regions exhibiting both types of alignments are considered collapsed relative to the other assembly, while those with only non-reciprocal best hit alignments are considered expanded (https://www.ncbi.nlm.nih.gov/genome/tools/remap/docs/alignments). Evaluating the lengths of collapsed and expanded sequence on the chromosomes in both assemblies, we observed that all GRCh37 chromosomes exhibit more collapse than their GRCh38 counterparts (Figure 2). The increased variant representation in GRCh38 is responsible for much of this, as GRCh38 alternate loci scaffolds are implicated in the alignments of the ten largest GRCh37 collapsed regions, as well the ten largest GRCh38 expanded regions (Supplemental Worksheet S3). To assess the relative collapse and expansion of the two assemblies independent of the alternate loci, we compared the alignments of the non-redundant collection of sequences comprising the chromosomes and unlocalized and unplaced scaffolds (primary assembly units). Consistent with the full assembly alignments, we find that nearly all GRCh37 chromosomes exhibit a greater degree of collapse and less expansion than their GRCh38 counterpart, and also observe a correspondence between the most collapsed GRCh37 and most expanded GRCh38 assembly regions (Figure 2). From these analyses, we find that not only does the GRCh38 assembly gain additional sequence representation through the addition of alternate loci, but the GRCh38 chromosomes provide more accurate representations of duplicated or paralogous regions than those of GRCh37.

**Figure 2.**
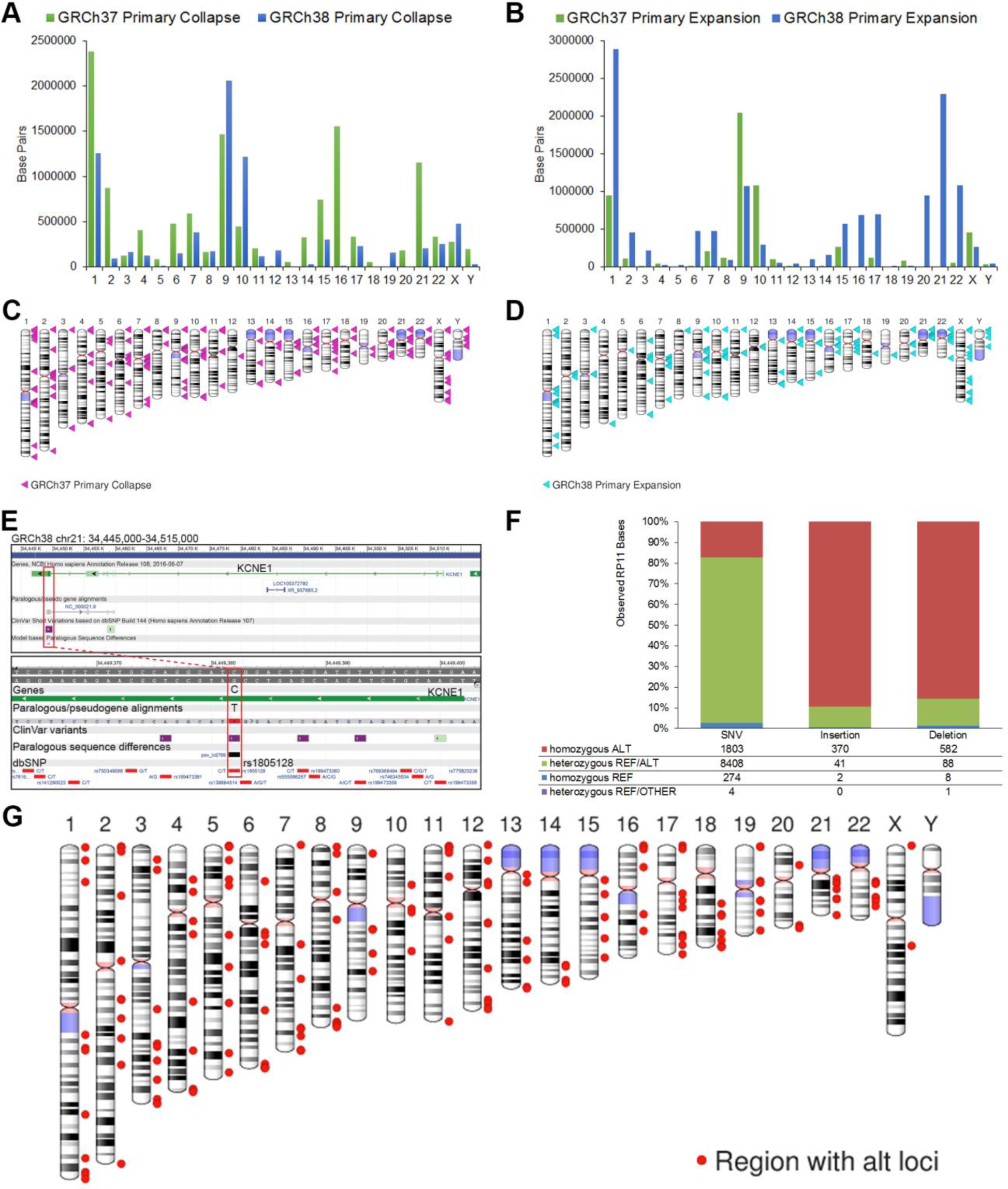
Evaluation of assembly updates. (A, B) Plots showing the per-chromosome lengths of sequence collapse (A) and expansion (B) of the GRCh37 (green) and GRCh38 (blue) primary assembly units (from which alt loci excluded), based on their assembly-assembly alignment. (C, D) Ideograms showing the location of collapsed sequence in GRCh37 and expanded sequence in GRCh38, based on the alignments of their primary assembly units. The sites are largely overlapping. (E) Browser view of *KCNE1* on GRCh38 chr 21. The lower panel shows a zoomed view of the top, illustrating a paralogous sequence alignment and paralogous variant (psv) overlapping SNP rs1805128, a putatively pathogenic ClinVar variant we observed remapping to multiple locations in GRCh38, due to the addition of paralogous sequence. Because prior assembly versions lack this paralog (red box), reads may map incorrectly in this region, and the pathogenicity of the variant and associated diagnostic calls should not be based only on such analyses. (F) Plot showing the allele distribution in RP11 WGS reads for the set of GRCh37 bases located in RP11 assembly components that were flagged as putative errors because they were not observed in the 1000 Genomes phase 1 dataset. (G) Ideogram showing the distribution of regions containing alternate loci scaffolds in GRCh38.

To assess the implications of these expanded sequences, we examined their effect at GRCh37 and GRCh38 genomic sites annotated with the subset of dbSNP Build 147 variations described in ClinVar (Landrum et al. 2014). In one analysis, we aligned reads from the Ashkenazi female sample NA24143 with BWA-MEM and evaluated ClinVar sites that have coverage with at least one MAPQ 20 or greater alignment in the GRCh37 and GRCh38 primary assemblies (Zook et al. 2016; Li 2013). We found variants annotated at 10 locations, representing 3 different chromosomes, which gained such coverage in GRCh38 (Table 4). Each of these regions was explicitly curated to remove redundant sequence or correct haplotype expansions in GRCh37. Variant calls missed on GRCh37 at these locations due to the artificial presence of confounding sequence should now be possible to call on GRCh38. We also identified variants annotated at 135 locations, associated with 6 different genomic regions, at which such coverage was lost in GRCh38. All are correlated with GRC curations in which allelic or paralogous sequence was added in GRCh38, suggesting that read alignments at these loci in GRCh37 may give rise to false variation calls. Together, these analyses show that assembly updates associated with the representation of duplicated or paralogous sequence affect read alignment, including at clinically relevant loci, which may have critical impacts on variant discovery and diagnosis. In a second analysis, we used the same collection of ClinVar variants to evaluate the impact of assembly updates on the remapping of data from GRCh37 to GRCh38. We identified a subset of unique GRCh37 ClinVar variants (n=210), including at least one described as putatively pathogenic, that mapped ambiguously to the GRCh38 primary assembly. These variants are associated with 9 genomic regions, all of which underwent deliberate curation to add sequence deemed missing from prior assembly versions (Table 4 and Supplemental Worksheet S4). In some instances, the newly added sequence exhibits paralogous variation and represents what was previously declared to be the non-reference allele (Figure 2). The results from this small survey of human variation further illustrate the potential impact that assembly updates can have on variant calling and diagnosis and demonstrate the importance of performing such evaluations on the GRCh38 assembly, with its expanded sequence representation.

**Table 4.**
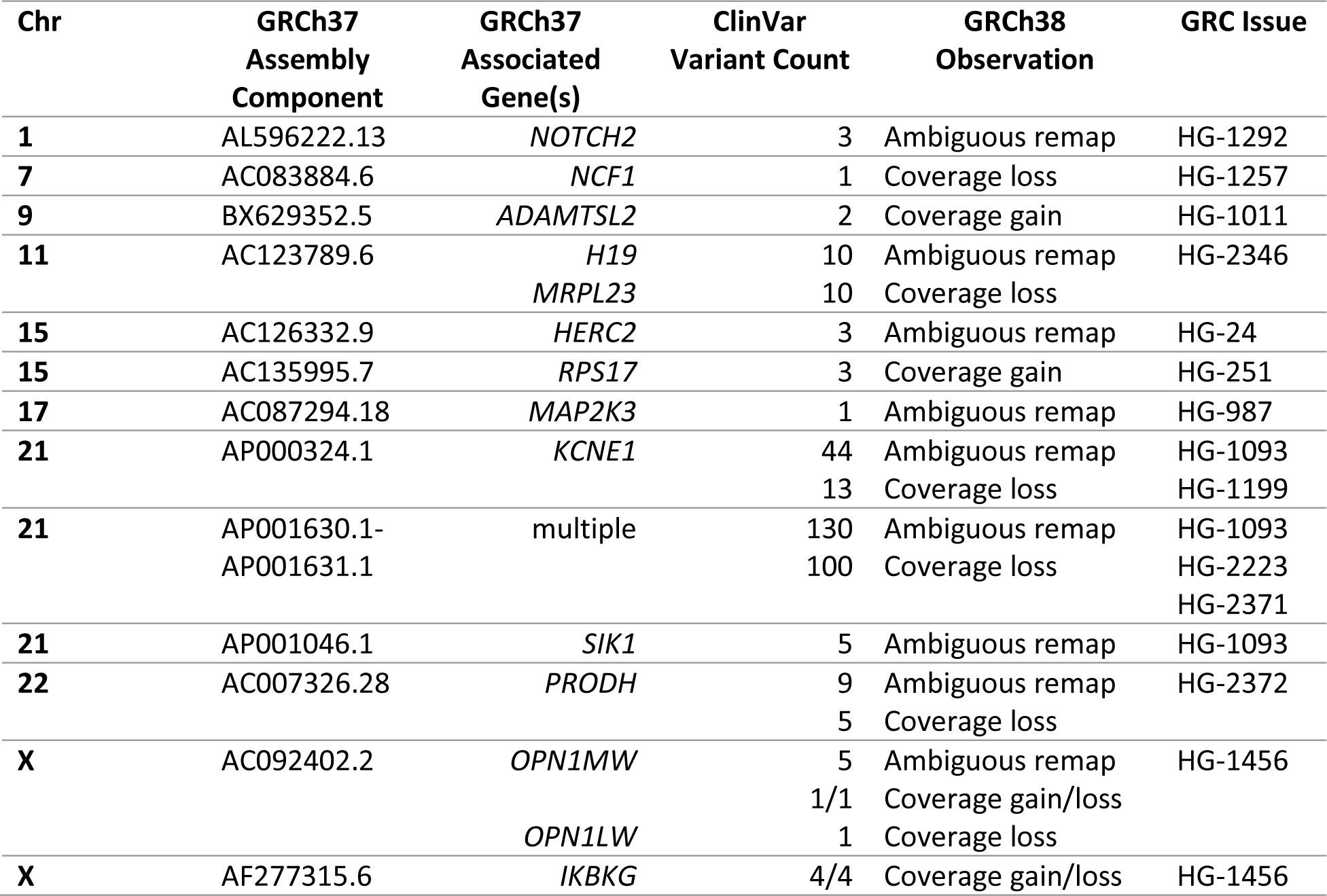
Evaluation of ClinVar variants

### Base Updates

In addition to large-scale curations, we also performed targeted sequence updates. Because erroneous reference bases, estimated to occur at a rate of 10^−5^, can result in incorrect variant calls, complicate gene annotation, and in the case of indels, complicate read alignments, we sought to identify and correct such sites (International Human Genome Sequencing Consortium 2004). We considered a set of 15,244 GRCh37 SNVs and 2375 indels with a minor allele frequency (MAF)=0 in the phase 1 analysis of the 1000 Genomes project or that could not be identified in a *k*-mer analysis as candidate reference errors (1000 Genomes Project Consortium et al. 2012, 2010) (Supplemental Methods). For the subset of sites located in RP11 BAC components, we sought to validate the assertion that the reference alleles represent errors. We examined allele distributions in the RP11 genome by aligning Illumina WGS reads from RP11 and looking for evidence of the reference base in the sample. Among the candidate sites, we observed that 80% of SNVs, 10% of insertions and 13% of deletions were heterozygous in RP11 (Figure 2), indicating that they were not reference errors. This analysis demonstrates the difficulty in distinguishing private or very low frequency alleles from error, even with large variation data sets. To ensure we retained the haplotype structure of the RP11 BAC components in the reference assembly, we did not update the observed RP11-derived heterozygous candidate sites in GRCh38. Given the admixed ancestry of the RP11 donor, it remains to be determined whether these otherwise unknown alleles are preferentially associated with a specific population background.

For the remaining sites, we used reads from samples in the 1000 Genomes phase 1 dataset or RP11 to generate short WGS contigs whose sequence overlapped the target site and surrounding bases (Supplemental Methods). We validated these “mini-contigs” by alignment to GRCh37, confirming that they differed only at the target site and contained the expected alternate allele and added them to the assembly. In a small number of cases, WGS contigs from other human assemblies or genomic PCR products were instead used to update bases. We updated an additional 376 sites identified during the course of other curation activity, that while not monomorphic, were either deemed universally rare according to 1000 Genomes phase 1 analysis or that had been reported by clinical testing labs and annotators to have a substantial negative impact on clinical variation analyses or annotation. In total, 8,249 sites were updated (Supplemental VCF S1, Supplemental VCF S2), 35 of which are annotated as ClinVar variants in GRCh37. These targeted updates represent the first large-scale effort to correct base-pair level errors in the reference.

### Alternate Loci Additions

In addition to adding sequence at assembly gaps and providing representation for missing copies of segmental duplication, we increased the number of alternate loci scaffolds to provide more representation for population variation in the reference. GRCh38 includes 261 scaffolds representing 178 genomic regions (Figure 2). As described previously, these alternate loci improve read mapping, provide the only reference representation for >150 genes, and capture sequence from the 1000 Genomes “decoy” used as a read sink for GRCh37, endowing it with chromosome context (Church et al. 2011, 2015; 1000 Genomes Project Consortium et al. 2015). Of particular note, GRCh38 includes 35 different representations for the immune-related Leukocyte Receptor Complex on chromosome 19 and two additional haplotype resolved paths of the highly variable and complex Spinal Muscular Atrophy (SMA) region on chromosome 5 (Schmutz et al. 2004; Pyo et al. 2010). The GRC website provides additional information about alternate loci with a series of region-specific pages that provide a graphical display and a report of associated curation issues (https://www.ncbi.nlm.nih.gov/projects/genome/assembly/grc/human/).

### Impacts on Read Mapping

We evaluated the impact of the cumulative set of GRCh38 updates on read mapping. Reads from the Ashkenazi sample used for the ClinVar analysis were aligned to the GRCh37 and GRCh38 primary assemblies and to the GRCh38 full assembly (Supplemental Material). While the GRCh37 primary assembly is an excellent mapping target, with 99.92% of reads aligned, we find that 64.32% of the unmapped reads are now mapped to the GRCh38 primary assembly. Consistent with the assembly curation effort, we observe many of these previously unmapped reads aligning to new sequences added at GRCh37 gaps (Figure 3). This demonstrates that the updates found on the GRCh38 reference assembly chromosomes make them a more robust substrate for analyses than the prior assembly version. We also find that 23.71% of reads that are still unmapped on the GRCh38 primary assembly map to the GRCh38 full assembly, which includes the alternate loci. We frequently observe these reads aligning to sequence unique to the alternate loci, validating GRC efforts to expand reference sequence representation with alternate loci (Supplemental Figure S1).

**Figure 3.**
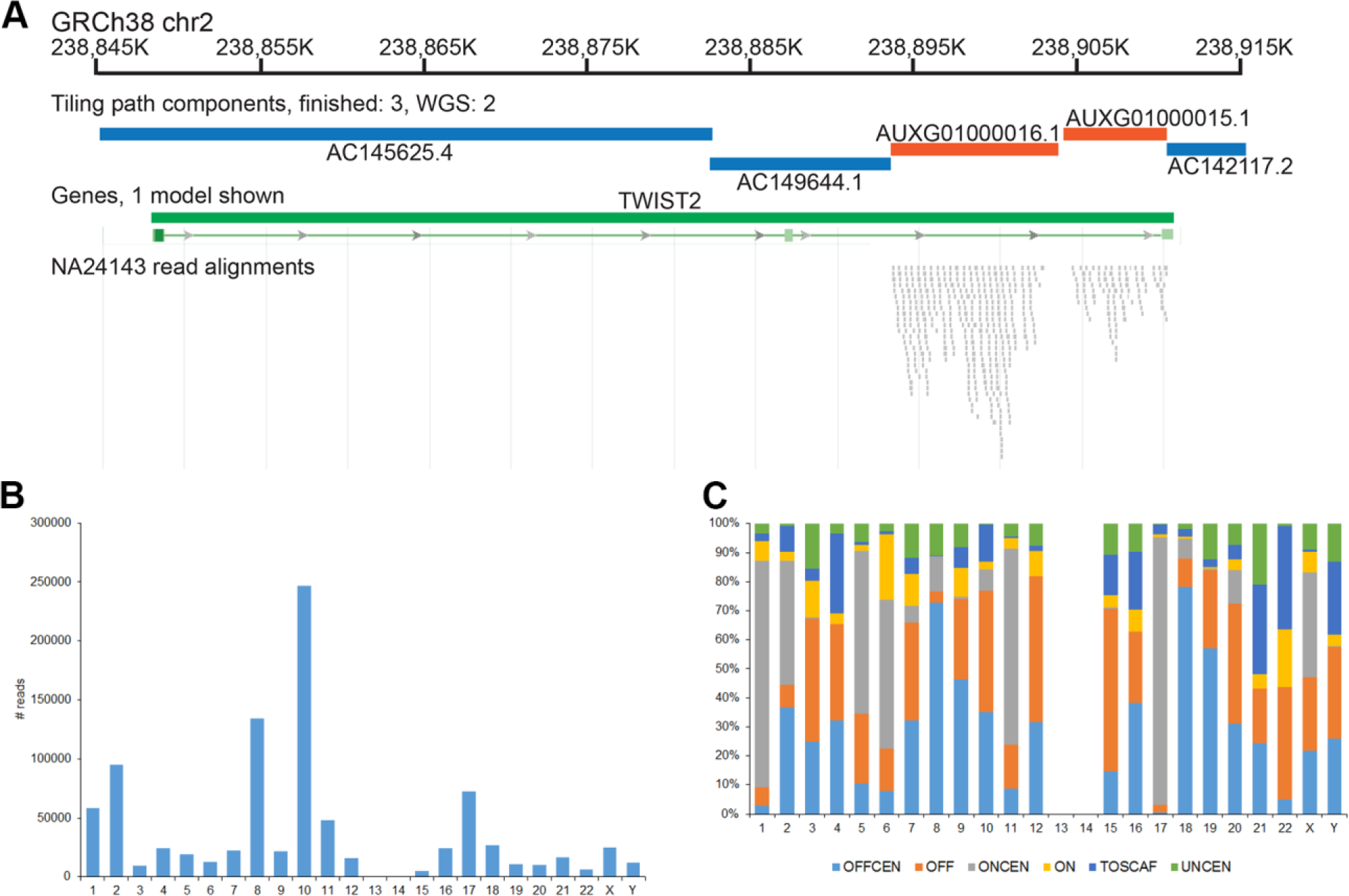
NA24143 read alignments to GRCh38. (A) Schematic showing the alignment of a subset of reads unmapped on GRCh37 to GRCh38. Reads align to GRCh38 at the position of components that were added to span a GRCh37 assembly gap (orange). (B) Graph showing counts of reads uniquely mapped to unchanged regions of GRCh37 that uniquely map to non-equivalent locations in GRCh38. (C) Chart describing the GRCh38 distribution of reads from Figure 3B, categorized by sequence location (same or different chromosome/scaffold) and sequence type (centromeric vs. non-centromeric). OFFCEN = movement to centromeric sequence on different chromosome; OFF = moment to non-centromeric sequence on different chromosome; ONCEN = movement to centromeric sequence on same chromosome; ON = movement to non-centromeric sequence on same chromosome; TOSCAF = movement to a non-centromeric unlocalized or unplaced scaffold; UNCEN = movement to a unplaced scaffold containing centromere-associated sequence.

While assembly updates are expected to alter read alignments in changed regions, we also investigated their impact on read mappings in the 2.6 Gbp of unchanged reference sequence. We find that 4.19% of read pairs that map uniquely, albeit imperfectly, to the GRCh37 primary assembly in an unchanged assembly region move to a new location with a different underlying assembly component in GRCh38. Approximately one-third of these moved pairs are also uniquely mapped to GRCh38 (Supplemental Table S2). We also analyzed the movement of individual reads from the moved pairs with respect to location (on- or off-chromosome) and sequence type (centromeric or non-centromeric). We find that both the extent and patterns of read movement are unique to each chromosome (Figure 3). Consistent with a non-random pattern of movement, we observe distinct pairings of assembly components over-represented as GRCh37 and GRCh38 mapping targets for each chromosome. Among reads belonging to moved pairs that also map uniquely to GRCh38, transitions to the modeled and unplaced GRCh38 centromere sequences predominate, but shifts to non-centromeric sequence still account for about 25% of total movement (Figure 3). Together, these analyses demonstrate that the assembly updates and alternate loci in GRCh38 not only make it a more complete mapping target, but that updates also exert an effect beyond their borders. As a result, we recommend use of GRCh38 for new genome-wide analyses in addition to studies specifically associated with changed regions.

### De novo assembly evaluations

The majority of reference assembly updates in GRCh38 used finished genomic clones. New reference-quality sequence sources are needed, though, as generation of finished sequence from clone libraries is in significant decline due to cost and some remaining assembly gaps occur in regions recalcitrant to cloning. A growing collection of human genomes in INSDC databases, a prerequisite for any sequence that will contribute to the reference assembly, that were sequenced and assembled with new technologies are candidates for use in assembly improvement (Pendleton et al. 2015; Zook et al. 2016; English et al. 2015; Bradnam et al. 2013; Earl et al. 2011; Vezzi et al. 2012). However, WGS assembly sequences have historically not been considered reference quality, raising concerns about their use in reference genome assembly curation. The essentially homozygous genomes of CHM1 and CHM13 have great potential for use in future updates due to the proven usefulness of haploid resources in resolving complex regions (Steinberg et al. 2014; Chaisson et al. 2015a; Huddleston et al. 2014; Berlin et al. 2015). We therefore generated the first collection of WGS de novo assemblies of CHM1 and CHM13 from two new sets of publicly available read data (SRP044331 and SRP051383), using both FALCON (Chin et al. 2016) and Celera Assembler (Berlin et al. 2015), with the intention of evaluating them with respect to reference assembly characteristics. We initially compared basic statistics for these assemblies to each other and to the GRCh38 assembly (Table 1). In addition, we compared the CHM1 assemblies to CHM1_1.1, a hybrid clone and short-read based reference guided assembly of CHM1 (Steinberg et al. 2014). We used Illumina data to determine the QV scores for each de novo assembly, providing a measure of base-pair level accuracy. All assemblies exhibit overall high quality, each with a QV near or >40. For both samples, we found that total lengths of the new assemblies were consistent with respect to one another and to GRCh38 or CHM1_1.1. The contig N50s of the new assemblies exhibited more variability, demonstrating that while all assemblies will have most of the same sequence for a given sample, they vary in how it is put together. Strikingly, even without scaffolding, many of these N50s are comparable with the scaffolded N50s of other recently published de novo WGS assemblies, auguring well for their ability to contribute to gap closure curation efforts (Pendleton et al. 2015; Chaisson et al. 2015a; Wang et al. 2008; Berlin et al. 2015).

To ensure continued reference quality, it is critical that gap-closing sequences or new alternate loci exhibit long-range accuracy. Thus, we used a number of other metrics to evaluate overall assembly quality. We performed paired end alignments of clones from a CHM1 derived BAC library (CHORI-17) to the CHM1 assemblies (Supplemental Table S3). We find that 96.64% (GCA_001307025.1) and 96.53% (GCA_001297185.1) of placements are unique and concordant, comparable to the 96.22% reported in an equivalent analysis of the CHM1_1.1 assembly (Steinberg et al. 2014). We also compared assembly contigs to BioNano optical maps we generated for the CHM1 and CHM13 assemblies (Table 5). For CHM1 and CHM13, total numbers of indels and inversions were very low and comparable among assemblies in each set, demonstrating the overall high quality of these assemblies. Scaffolding the contigs with the optical map data resulted in near doubling of the N50s and a 20-fold decrease in molecule number (Table 6). Based on these analyses, sequences from all of these assemblies appear to offer promise for use in gap closure or addition to the assembly as alternate loci.

**Table 5:**
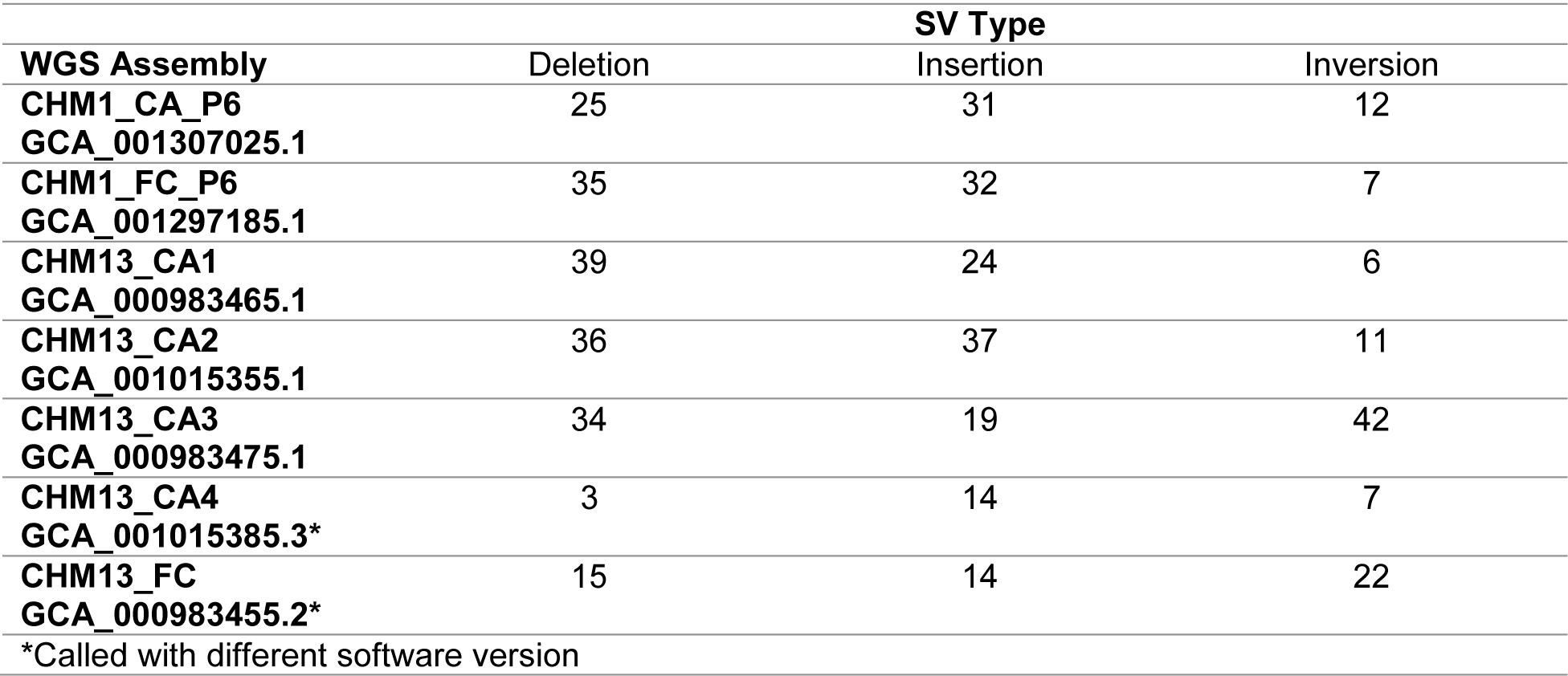
Structural Variation Calls in WGS-BioNano Assembly Comparison

| WGS Assembly | SV Type |  |  |
| --- | --- | --- | --- |
|  | Deletion | Insertion | Inversion |
| **CHM1_CA_P6**<br>**GCA_001307025.1** | 25 | 31 | 12 |
| **CHM1_FC_P6**<br>**GCA_001297185.1** | 35 | 32 | 7 |
| **CHM13_CA1**<br>**GCA_000983465.1** | 39 | 24 | 6 |
| **CHM13_CA2**<br>**GCA_001015355.1** | 36 | 37 | 11 |
| **CHM13_CA3**<br>**GCA_000983475.1** | 34 | 19 | 42 |
| **CHM13_CA4**<br>**GCA_001015385.3\*** | 3 | 14 | 7 |
| **CHM13_FC**<br>**GCA_000983455.2\*** | 15 | 14 | 22 |
| *Called with different software version |  |  |  |

**Table 6:** Hybrid Assembly Statistics

|  | BioNano<br>Assembly<br># Contigs | Hybrid<br>Assembly<br># Scaffolds | Hybrid<br>Assembly<br>Scaffold N50<br>(Mb) | Hybrid<br>Assembly<br>Total Size<br>(Gb) | WGS<br>Assembly<br>Conflicts | BioNano<br>Assembly<br>Conflicts |
| --- | --- | --- | --- | --- | --- | --- |
| CHM1_CA_P6<br>GCA_001307025.1 | 2473 | 221 | 40.04 | 2.82 | 52 | 63 |
| CHM1_FC_P6<br>GCA_001297185.1 | 2473 | 161 | 47.60 | 2.84 | 45 | 51 |
| CHM13_CA1<br>GCA_000983465.1 | 3593 | 238 | 29.46 | 2.81 | 39 | 48 |
| CHM13_CA2<br>GCA_001015355.1 | 3593 | 229 | 35.09 | 2.82 | 56 | 83 |
| CHM13_CA3<br>GCA_000983475.1 | 3593 | 379 | 14.80 | 2.82 | 29 | 31 |
| CHM13_CA4*<br>GCA_001015385.3 | 3593 | 258 | 27.83 | 2.82 | 43 | 55 |
| CHM13_FC*<br>GCA_000983455.2 | 3593 | 229 | 21.66 | 2.83 | 63 | 84 |
| *Conflicts called with different software version than other assemblies (see Methods) |  |  |  |  |  |  |

We further evaluated assembly quality with feature response curves (FRC) generated with mapped Illumina read pairs as input to FRC^bam^ (Figure 4) (Vezzi et al. 2012). Even though N50s differ by more than a factor of two among the assemblies, all FRC scores are high and comparable, indicating their overall quality and that additional joins in assemblies with longer N50s do not introduce significant error. However, because repetitive sequences have typically been prone to collapse in WGS assemblies, we also used FRC curves to evaluate compression and expansion in each of the assemblies. Once again, we see that all assemblies fared well with respect to this metric, clustered at the center, with only minor differences between assemblers or parameters for a given sample. The long reads and lack of allelic variation in these new assemblies likely underlie these observations (Huddleston et al. 2014).

**Figure 4.**
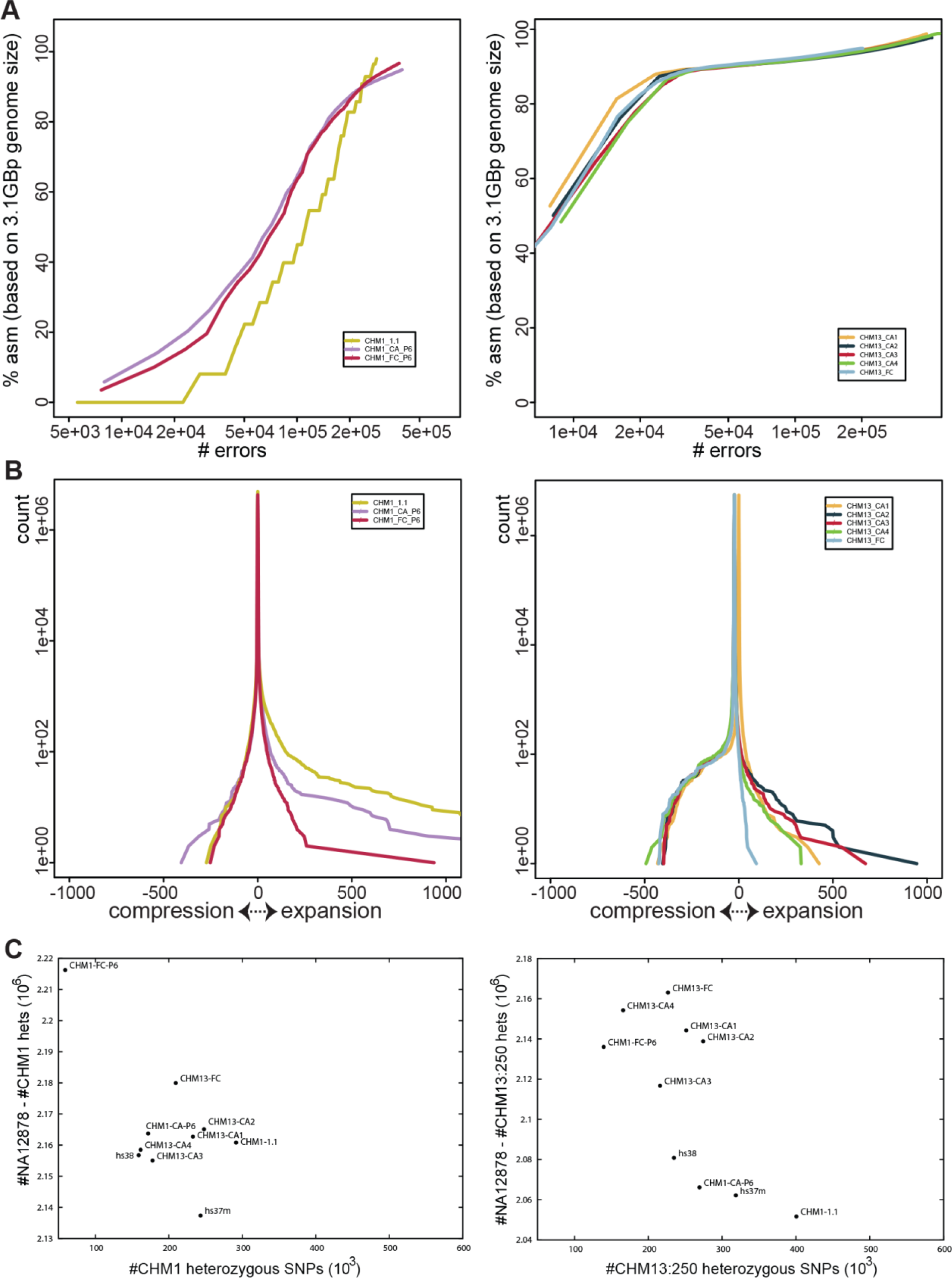
Evaluation of CHM1 and CHM13 assemblies. (A) FRC error curve for CHM1 (left) and CHM13 (right) assemblies. CHM1_1.1 is provided for comparison with the CHM1 de novo assemblies. The x-axis is log-scaled. (B) FRC compression-expansion curve for CHM1 (left) and CHM13 (right) showing distribution of mapped reads. Divergence from the center indicates compression (negative) and expansion (positive). (C) Heterozygous SNPs called on the CHM1 and CHM13 de novo assemblies, CHM1_1.1 and GRCh38 using NA12878 and CHM1 (left) and CHM13 (right) aligned FermiKit assemblies. The x-axis represents potential false positives, while the y-axis measures potential true positives; optimal assemblies appear in the upper left of the plot.

We also appraised the assemblies by variant calling with FermiKit, in which heterozygous variant calls based on alignment of haploid samples are considered false positives, likely caused by assembly collapse of tandem repeats and/or segmental duplications (Figure 4, Supplemental Materials, Supplemental Figure S2) (Li 2015, 2014). Heterozygous calls on the collections of CHM1 and CHM13 assemblies were measured using three different haploid de novo assemblies, and evaluated with respect to heterozygous calls from the diploid NA12878 sample. These analyses uniformly show that for the CHM1 sample, the FALCON-based assembly is a better substrate for variant calling, but also suggest that Celera Assembler produces a better variant calling substrate for the CHM13 sample. Comparison to GRCh37 and GRCh38 suggests that these new haploid assemblies may serve as more reliable substrates for variant calling than the reference assembly, though further analysis is needed to determine whether improvements occur in genomic regions of interest. However, as variant calling is only one use case for the reference assembly, we also examined other facets of these de novo assemblies.

Gene content is another important metric for assembly quality, especially if the assembly will be used as an annotation substrate. We examined three aspects of RefSeq transcript alignments to the CHM1 and CHM13 assemblies to assess different aspects of assembly quality. Total gene representation reflects overall assembly quality and content, co-placement of genes reflects collapsed segmental duplications, while frameshift analysis provides information about the accuracy of gene representation within the assembly (Table 7). We find that all assemblies compare favorably to each other and to GRCh38 with respect to total content of gene representation. In contrast, we find that all CHM1 and CHM13 assemblies exhibit a substantially greater number of transcripts that are dropped due to conflicting placement with transcripts representing other genes, compared both to the GRCh38 reference assembly and to the CHM1_1.1 assembly (Table 7). The genes associated with co-placed transcripts are largely shared within and between assemblies derived from CHM1 or CHM13 and are dominated by paralogous genes, many of which reside in multi-megabase highly complex and/or segmentally duplicated regions (Supplemental Worksheet S5, Supplemental_GFF3_S1.tar.gz, Supplemental Worksheet S6, Supplemental_GFF3_S2.tar.gz). The genomic locations associated with the transcripts on these lists may reflect regions still recalcitrant to assembly with current read lengths and algorithms. These lists also include haplotype-specific or copy-number variant genes, for which co-placement occurs when they are absent from the sample haplotype. In contrast to the GRCh38 reference assembly, in which alternate loci provide representation for multiple haplotypes at many loci, the CHM1 and CHM13 samples represent only a single haplotype and are expected to have a slightly lower overall gene content, which may also contribute to the higher number of co-placed genes on these assemblies relative to GRCh38. However, there are 35-40% fewer transcripts dropped from the CHM1_1.1 assembly due to co-placement than from the FALCON or Celera Assembler CHM1 assemblies, indicating that assembly method has a substantial impact on gene representation. In the context of reference assemblies, these findings demonstrate that caution is required when using assemblies that have been deemed “high quality”. Gene content must be considered as part of the determination of whether an assembly is suitable for use as a reference or in reference curation. Assembly method can have a striking impact on the accuracy of predicted proteins, as can sequencing technology (Florea et al. 2011). To assess the quality of protein representation in these assemblies, we identified RefSeq alignments containing frameshifting (FS) indels in coding sequence. We observe that the number of transcripts aligning with frameshifting indels is much higher in these new assemblies compared to GRCh38 or CHM1_1.1 (Table 7). Additionally, for both samples, we find that the likelihood of a FS protein being unique to a particular assembly or shared among all assemblies is roughly equivalent, further confirming the influence of assembly method on protein prediction. Using the subset of FS proteins not common to all assemblies as a denominator, we examined the percentage of uniquely FS proteins in each assembly. For the CHM13 sample, an average of 50% of FS proteins were unique to each assembly, ranging from a high of 61% in the FALCON assembly, to a low of 40% in the Celera Assembler assemblies. For CHM1, both assembly methods performed similarly, with ~50% of FS proteins unique to either assembly. We also looked at the subset of FS proteins common to all de novo assemblies for each sample, which are most likely to represent true variation, and/or arise from issues with the read data or genomic regions problematic for all assembly methods. Consistent with the former, we find that the *GRIN3B* gene has a frameshifting indel in all CHM1 and CHM13 assemblies that corresponds to rs10666583, a known inactivating variation associated with susceptibility to schizophrenia (Matsuno et al. 2015). While further analyses are required to understand the differences at the assembly sequence level and to assess the effect that assembly polishing tools such as Pilon might have (Walker et al. 2014), these data clearly demonstrate the variability in gene representation that can arise due to assembly method. Together, our analyses indicate that recent long read assemblies have good continuity, a low error rate, and a high rate of gene completeness compared to previous de novo efforts. They should prove valuable for resolving a subset of remaining reference assembly gaps and providing variant sequence representations. However, the reference still provides better representations of long repeat structures and genes, meaning that further advances in assembly methods are needed if WGS assemblies are truly to be recognized as reference quality genomes.

**Table 7.** RefSeq evaluation of de novo assemblies

| Assembly | Not Aligned (%) | Split Alignment (%) | Coverage <95% (%) | Dropped coding transcripts | Dropped non-coding transcripts | Proteins with frameshifts* |
| --- | --- | --- | --- | --- | --- | --- |
| **GRCh38**<br>**GCA_000001405.15** | 22 (0.04%) | 10 (0.02%) | 17 (0.04%) | 2 | 0 | 19 |
| **CHM1_1.1**<br>**GCA_000306695.2** | 89 (0.17%) | 40 (0.08%) | 257 (0.65%) | 171 | 123 | 218/221 |
| **CHM1_CA_P6**<br>**GCA_001307025.1** | 117 (0.23%) | 291 (0.23%) | 426 (1.08%) | 226 | 160 | 983 |
| **CHM1_FC_P6**<br>**GCA_001297185.1** | 65 (0.13%) | 171 (0.34%) | 234 (0.60%) | 214 | 167 | 1012 |
| **CHM13_CA1**<br>**GCA_000983465.1** | 50 (0.10%) | 345 (0.68%) | 386 (0.98%) | 274 | 213 | 503 |
| **CHM13_CA2**<br>**GCA_001015355.1** | 49 (0.10%) | 320 (0.63%) | 335 (0.85%) | 272 | 213 | 439 |
| **CHM13_CA3**<br>**GCA_000983475.1** | 46 (0.09%) | 616 (1.22%) | 632 (1.61%) | 240 | 187 | 627 |
| **CHM13_CA4**<br>**GCA_001015385.3** | 50 (0.10%) | 400 (0.79%) | 404 (1.03%) | 259 | 197 | 450 |
| **CHM13_FC**<br>**GCA_000983455.2** | 94 (0.18%) | 482 (0.96%) | 568 (1.44%) | 281 | 202 | 346 |
| **50867 RefSeq transcripts were aligned to each assembly** |  |  |  |  |  |  |
| **\*GRCh38 frameshifts exclude alternate loci** |  |  |  |  |  |  |

## Discussion

The human reference genome assembly, initially released over a decade ago, remains at the nexus of basic and clinical research. Like the continually changing landscape in which it exists, the reference assembly also evolves. As we have described, GRCh38, the current version of this resource, exhibits improved assembly statistics, contains corrected representations of several large-scale clinically relevant regions and provides new sequence content. This content both captures previously missing genomic sequence and provides representations of population genomic diversity. The updates to the assembly render it an improved annotation substrate and alter its characteristics as a mapping target. Together, the suite of changes introduced in GRCh38 make it the most complete and accurate representation of the human genome yet produced and we recommend its use over prior assembly versions for all types of analysis.

In order to establish the relevancy of a clone-based reference assembly in the context of new sequencing and assembly technologies, we also generated and evaluated several de novo long read based assemblies representing the CHM1 and CHM13 haploid genomes with respect to each other and GRCh38. All proved to be high quality and demonstrate the capabilities of FALCON and Celera Assembler to generate robust assemblies from large scale, complex genomic data sets. Nonetheless, each assembly method imparted distinct characteristics to the haploid assemblies, and none could be considered the best genome representation by all metrics evaluated. We suggest that de novo assemblies may be further improved by development to support the use of additional datasets, such as Illumina reads or genomic clones, as input to the assembly process, or by post-processing with various error correcting tools. The de novo assemblies also demonstrate the challenges and limitations in transforming data associated with repetitive or complex genomic regions from a rich graph-based assembler representation to a narrower linear assembly representation. It may be desirable to adjust parameters to convey different aspects of the data, such as length, variation content or sequence quality, in order to produce assemblies best suited to different types of analyses. Notably, such suites of sequence representations could be captured in the current reference assembly model as alternate loci scaffolds, and de novo assemblies may further contribute to the reference in this way.

Our analysis of GRCh38 and the de novo assemblies demonstrates that the reference assembly remains the most comprehensive and highest quality representation of the human genome, capable of supporting the widest range of analyses and discoveries. However, we also foresee an evolving role for the reference genome assembly in the context of two anticipated sea-changes in genome biology that will be realized by ongoing development for technological and computational methods: (1) a proliferation of reference-quality individual diploid genome assemblies and (2) a comprehensive graph-based representation of genome-wide population variation. In both contexts, the reference assembly is likely to serve as a point of integration. In an era of personalized medicine, we anticipate the integration of data analyses performed on individual genomes through the reference assembly. Regardless of its quality, an assembly representing an individual genome will be limited in its representation of variation. The reference assembly provides context for both the scale and types of variation that will be observed from one sample to the next. Using the reference in this role presents a mechanism for transferring individual interpretations to populations. However, these efforts will require tools and resources for comparative analysis. Without continued development in this area, the challenges incurred today in evaluating analyses performed on different versions of the reference assembly, or transitioning datasets between them, will persist and be magnified as the extent of the differences between individuals will be considerably greater than those between reference assembly versions. GRCh38, with its robust genome representation and well-characterized assembly features provides the framework for this development.

The reference is also a framework for the establishment of a genome graph that represents population variation. This is a natural step in the evolution of the scientific role of the reference genome assembly. Conceived from the outset as a model of the human pan-genome, the current reference now contains not only chromosome sequences depicting a mosaic of haplotypes from different individuals, but includes alternate loci scaffolds that provide multi-allelic and -haplotypic representation for regions across the genome. Because the alignments that define the relationship of these scaffolds to the chromosomes are integral pieces of the assembly model, we submit that the reference has already started the transition into a graph-based depiction of the human genome. As genome graphs progress further into nonlinear forms, the reference chromosome sequences are well-suited to serve as a central path against which variation is described or annotations are made, while the alternate loci provide a subset of high-quality and curated branches (Paten et al. 2014; Nguyen et al. 2015; Dilthey et al. 2015; Novak et al. 2016). The Global Alliance for Genomic Health (GA4GH) are using the GRCh38 assembly with alternate loci in a pilot graph-building project (https://github.com/ga4gh/schemas/wiki/Human-Genome-Variation-Map-(HGVM)-Pilot-Project). Ongoing reference curation efforts are aimed at providing additional representations for genomic diversity and have added more than 40 novel patches since the initial release of GRCh38. The continued improvement of the reference assembly does therefore not put it in conflict with these new models, but instead will serve to improve them, as it provides a more robust representation of the sequences and relationships that they will portray.

In an idealized view, the reference assembly should be improved until this critical resource is sufficiently complete that it: (1) provides chromosome context for any identified human sequence of 500 bp or greater (Church et al. 2011), (2) enables unambiguous data interpretation at all clinically relevant loci and (3) introduces no systematic error or bias in genome-wide analyses. The substantial improvements and changes represented in GRCh38 move us closer to this ideal on all three points. The analyses of the high quality de novo haploid CHM1 and CHM13 assemblies show that there may soon be new resources that will bring us even nearer to this goal. However, the challenges in migrating datasets and paucity of tools for working with allelic sequence representations such as alternate loci and patch scaffolds presents a barrier to the adoption of new assemblies, despite their improvement over prior versions (Church et al. 2015). Our ability to address these challenges will, in part, define the point at which the reference representation is deemed sufficient on all three goals to render further improvements unwarranted. As the community of reference assembly users draws ever closer to that point, we caution that we must let the biology, rather than the technology or an abstracted goal, be the primary driver for that decision. In keeping with that view, we foresee a continued need for assembly evaluation in the context of the ever-evolving landscape of genome research.

## Methods

### Transcript Evaluation of Assemblies

Alignments were performed and analyzed as described in the Supplementary Methods of (Shi et al. 2016). However, in contrast to the RefSeq transcripts, we evaluated coverage for the GENCODE data over the full transcript, rather than the CDS, because we did not have the CDS information.

### Assembly-assembly alignments

Assemblies were aligned using software version 1.7 of the NCBI pipeline as described in the Methods of (Steinberg et al. 2014). The alignments and alignment reports are available from the NCBI Remap FTP site: ftp://ftp.ncbi.nlm.nih.gov/pub/remap/Homo_sapiens/1.7/ (Kitts et al. 2016). We evaluated chromosome-level collapse and expansion in these alignments with https://github.com/deannachurch/assembly_alignment/.

### ClinVar variant coverage analysis

We assessed coverage using the GATK DepthOfCoverage tool (McKenna et al. 2010), with the parameter --minMappingQuality 20.

We used the following VCF files containing ClinVar variants on the GRCh37 and GRCh38 assemblies to define the sites at which to assess coverage: ftp://ftp.ncbi.nlm.nih.gov/pub/clinvar/vcf_GRCh37/clinvar_20160502.vcf.gz ftp://ftp.ncbi.nlm.nih.gov/pub/clinvar/vcf_GRCh38/clinvar_20160502.vcf.gz

We measured the coverage for Illumina reads from sample NA24143 aligned to the GRCh37 and GRCh38 primary assembly units (described below) at these sites. Sites with zero coverage in GRCh37 were remapped to GRCh38 using the NCBI remapping service with default parameters (Kitts et al. 2016) (https://www.ncbi.nlm.nih.gov/genome/tools/remap/docs/api) and coverage re-evaluated. Sites with zero coverage in GRCh38 were remapped to GRCh37, and those with coverage were evaluated.

### ClinVar remapping analysis

We used the NCBI remapping service, with default parameters to remap the following variants from GRCh37 (GCF_000001405.13) to GRCh38 (GCF_000001405.26): ftp://ftp.ncbi.nlm.nih.gov/pub/clinvar/vcf_GRCh37/clinvar_20160502.vcf.gz

We manually reviewed the subset of variants with multiple re-mappings in the primary assembly unit.

### Base Updates

#### Evaluation of Candidate Bases in RP11 Assembly Components

We validated candidate erroneous bases in RP11 components with a pile-up analysis of the alignments of RP11 Illumina reads to GRCh37 in SRA run SRR834589.

Pile-up version: sra-pileup.2.3.2.11 (http://ncbi.github.io/sra-tools/), with the parameter --minmapq 20..

We used a cut-off of 90% to define homozygous and heterozygous reference and alternate allele calls at SNVs and a cut-off of 70% for indels. For indels, all non-homozygous alternate allele calls were manually reviewed. For SNVs, we manually reviewed all sites in which more than 2 alleles were called or in which alleles not expected for the corresponding dbSNP variant were reported.

### WGS mini-contig generation

Software:cortex_con_beta_0.04c (http://cortexassembler.sourceforge.net/). For additional details, see Supplemental Methods.

### Alignment of Illumina reads

2 × 150 bp reads from Ashkenazim trio sample NA24143 generated as described in (Zook et al. 2016) were aligned with BWA-MEM to the GRCh37 and GRCh38 assemblies. For additional details, see Supplemental Methods.

### CHM1/CHM13 Assembly Generation

Assemblies were either generated with Celera Assembler 8.3rc2 (Berlin et al. 2015) or with FALCON, an assembler based on HGAP (Chin et al. 2013, 2016),. For additional assembly details, see Supplemental Methods.

### Clone Placements

CH17 clone placements were performed and evaluated as described in (Steinberg et al. 2014; Schneider et al. 2013). On the GCA_001307025.1 assembly, the average insert length = 208,547 and the standard deviation = 19,641. On the GCA_001297185.1 assembly, the average insert length = 208, 596 and the standard deviation = 19,718.

### BioNano optical maps

Long CHM1 molecules were nicked and labeled according to the BioNano Genomics IrysPrep protocol and loaded on the IrysChip for genome mapping on the BioNano Genomics Irys System imaging instrument. Image detection, assembly and genome map alignment were performed using BioNano Genomics IrysSolve software tools. Each of the PacBio sequence assemblies were nicked in silico with BspQI to produce a cmap file, which reports the start and end coordinates and the placement of labels for each contig. BioNano Genomics software tools were then used to align each of the sequence assemblies to the CHM1 or CHM13 genome map, and SV detection software was run to generate the SV and hybrid stats provided in this paper (Supplemental Material).

### De novo assembly evaluation with Illumina read data

SRA accessions for reads used as input to Illumina read-based analyses (QV, FRC^bam^, FermiKit):

- CHM1: SRR2842672 (FRC), SRR642636-SRR642641 (FermiKit)
- CHM13-125: SRR2088062 and SRR2088063
- CHM13-250: SRR1997411
- NA12878: ERR194147 (FermiKit)

For additional details of these analyses, see Supplemental Methods.

## Data Access

All assemblies have been deposited in GenBank with the following accessions: (GRCh38) GCA_000001405.15, (WGS assemblies) GCA_001307025.1, GCA_001297185.1, GCA_000983465.1, GCA_001015355.1, GCA_000983475.1, GCA_001015385.3, and GCA_000983455.2. The corresponding read data for the WGS assemblies is in the SRA with the following accessions: SRP044331 and SRP051383.

## Acknowledgements

This research was supported by the Intramural Research Program of the NIH, National Library of Medicine, the Wellcome Trust (grant numbers WT095908, WT098051 and WT104947/Z/14/Z) and the European Molecular Biology Laboratory. SK and AMP were supported by the Intramural Research Program of the National Human Genome Research Institute, National Institutes of Health. This study utilized the computational resources of the Biowulf system at the National Institutes of Health, Bethesda, MD (http://biowulf.nih.gov). Work at MGI was supported by NIH grants 5U54HG003079 and 5U41HG007635.

The GRC wishes to acknowledge the invaluable assistance and contributions of the many external collaborators and the RefSeq and HAVANA annotation groups who shared data, expertise and advice in the effort to update the reference assembly sequence. A list of genome-wide and region-specific collaborators can be found at the GRC website: https://www.ncbi.nlm.nih.gov/projects/genome/assembly/grc/credits.shtml. Additionally, the GRC would like to thank Jim Knight and Stephan Schuster for submitting sequence from an RP11-based WGS assembly to INSDC (GCA_000442295.1), making it available for use in reference curation. The MGI would like to thank Susie Rock and Aye Wollam for their oversight of the assembly curation and finishing pipelines at the McDonnell Genome Institute. WTSI thanks Paul Heath, Guy Griffiths, Britt Killian and Eduardus Zuiderwijk for their technical and computational contributions. NCBI thanks Tayebeh Rezaie-Jami, Eugene Yaschenko, Avi Kimchi, and Karen Clark for their helpful discussion and expertise in content and data management. EBI thanks Bronwen Aken, Carlos García Girón and Amonida Zadissa.

## Disclosure Declaration

Richard Durbin: Dovetail Genomics

Deanna Church: 10X Genomics

Chen-Shan Chin: Pacific Biosciences

Paul Flicek: Omicia, Inc.

## References

1000 Genomes Project Consortium, Abecasis GR, Altshuler D, Auton A, Brooks LD, Durbin RM, Gibbs RA, Hurles ME, McVean GA. 2010. A map of human genome variation from population-scale sequencing. Nature 467: 1061–1073.

1000 Genomes Project Consortium, Abecasis GR, Auton A, Brooks LD, DePristo MA, Durbin RM, Handsaker RE, Kang HM, Marth GT, McVean GA. 2012. An integrated map of genetic variation from 1,092 human genomes. Nature 491: 56–65.

1000 Genomes Project Consortium, Auton A, Brooks LD, Durbin RM, Garrison EP, Kang HM, Korbel JO, Marchini JL, McCarthy S, McVean GA, et al. 2015. A global reference for human genetic variation. Nature 526: 68–74.

Antonacci F, Dennis MY, Huddleston J, Sudmant PH, Steinberg KM, Rosenfeld JA, Miroballo M, Graves TA, Vives L, Malig M, et al. 2014. Palindromic GOLGA8 core duplicons promote chromosome 15q13.3 microdeletion and evolutionary instability. Nat Genet 46: 1293–1302.

Bailey JA, Yavor AM, Massa HF, Trask BJ, Eichler EE. 2001. Segmental duplications: organization and impact within the current human genome project assembly. Genome Res 11: 1005–1017.

Berlin K, Koren S, Chin C-S, Drake JP, Landolin JM, Phillippy AM. 2015. Assembling large genomes with single-molecule sequencing and locality-sensitive hashing. Nat Biotechnol 33: 623–630.

Bradnam KR, Fass JN, Alexandrov A, Baranay P, Bechner M, Birol I, Boisvert S, Chapman JA, Chapuis G, Chikhi R, et al. 2013. Assemblathon 2: evaluating de novo methods of genome assembly in three vertebrate species. Gigascience 2: 10.

Cao H, Wu H, Luo R, Huang S, Sun Y, Tong X, Xie Y, Liu B, Yang H, Zheng H, et al. 2015. De novo assembly of a haplotype-resolved human genome. Nat Biotechnol 33: 617–622.

Chaisson MJP, Huddleston J, Dennis MY, Sudmant PH, Malig M, Hormozdiari F, Antonacci F, Surti U, Sandstrom R, Boitano M, et al. 2015a. Resolving the complexity of the human genome using single-molecule sequencing. Nature 517: 608–611.

Chaisson MJP, Wilson RK, Eichler EE. 2015b. Genetic variation and the de novo assembly of human genomes. Nat Rev Genet 16: 627–640.

Chin C-S, Alexander DH, Marks P, Klammer AA, Drake J, Heiner C, Clum A, Copeland A, Huddleston J, Eichler EE, et al. 2013. Nonhybrid, finished microbial genome assemblies from long-read SMRT sequencing data. Nat Methods 10: 563–569.

Chin C-S, Peluso P, Sedlazeck FJ, Nattestad M, Concepcion GT, Clum A, Dunn C, O’Malley R, Figueroa-Balderas R, Morales-Cruz A, et al. 2016. Phased Diploid Genome Assembly with Single Molecule Real-Time Sequencing. bioRxiv 056887. http://biorxiv.org/content/early/2016/06/03/056887.abstract (Accessed July 9, 2016).

Church DM, Schneider VA, Graves T, Auger K, Cunningham F, Bouk N, Chen HC, Agarwala R, McLaren WM, Ritchie GR, et al. 2011. Modernizing reference genome assemblies. PLoS Biol 9: e1001091.

Church DM, Schneider VA, Steinberg KM, Schatz MC, Quinlan AR, Chin C-S, Kitts PA, Aken B, Marth GT, Hoffman MM, et al. 2015. Extending reference assembly models. Genome Biol 16: 13.

Dennis MY, Nuttle X, Sudmant PH, Antonacci F, Graves TA, Nefedov M, Rosenfeld JA, Sajjadian S, Malig M, Kotkiewicz H, et al. 2012. Evolution of human-specific neural SRGAP2 genes by incomplete segmental duplication. Cell 149: 912–922.

Dilthey A, Cox C, Iqbal Z, Nelson MR, McVean G. 2015. Improved genome inference in the MHC using a population reference graph. Nat Genet 47: 682–688.

Earl D, Bradnam K, St John J, Darling A, Lin D, Fass J, Yu HOK, Buffalo V, Zerbino DR, Diekhans M, et al. 2011. Assemblathon 1: a competitive assessment of de novo short read assembly methods. Genome Res 21: 2224–2241.

English AC, Salerno WJ, Hampton OA, Gonzaga-Jauregui C, Ambreth S, Ritter DI, Beck CR, Davis CF, Dahdouli M, Ma S, et al. 2015. Assessing structural variation in a personal genome-towards a human reference diploid genome. BMC Genomics 16: 286.

Falconer E, Hills M, Naumann U, Poon SSS, Chavez EA, Sanders AD, Zhao Y, Hirst M, Lansdorp PM. 2012. DNA template strand sequencing of single-cells maps genomic rearrangements at high resolution. Nat Methods 9: 1107–1112.

Fan J-B, Surti U, Taillon-Miller P, Hsie L, Kennedy GC, Hoffner L, Ryder T, Mutch DG, Kwok PY. 2002. Paternal origins of complete hydatidiform moles proven by whole genome single-nucleotide polymorphism haplotyping. Genomics 79: 58–62.

Florea L, Souvorov A, Kalbfleisch TS, Salzberg SL. 2011. Genome assembly has a major impact on gene content: a comparison of annotation in two Bos taurus assemblies. PLoS One 6: e21400.

Genovese G, Handsaker RE, Li H, Altemose N, Lindgren AM, Chambert K, Pasaniuc B, Price AL, Reich D, Morton CC, et al. 2013a. Using population admixture to help complete maps of the human genome. Nat Genet 45: 406–414, 414e1–2.

Genovese G, Handsaker RE, Li H, Kenny EE, McCarroll SA. 2013b. Mapping the human reference genome’s missing sequence by three-way admixture in Latino genomes. Am J Hum Genet 93: 411–421.

Green RE, Krause J, Briggs AW, Maricic T, Stenzel U, Kircher M, Patterson N, Li H, Zhai W, Fritz MH-Y, et al. 2010. A draft sequence of the Neandertal genome. Science 328: 710–722.

Harrow J, Frankish A, Gonzalez JM, Tapanari E, Diekhans M, Kokocinski F, Aken BL, Barrell D, Zadissa A, Searle S, et al. 2012. GENCODE: the reference human genome annotation for The ENCODE Project. Genome Res 22: 1760–1774.

Hickey G, Paten B, Earl D, Zerbino D, Haussler D. 2013. HAL: a hierarchical format for storing and analyzing multiple genome alignments. Bioinformatics 29: 1341–1342.

Horton R, Gibson R, Coggill P, Miretti M, Allcock RJ, Almeida J, Forbes S, Gilbert JGR, Halls K, Harrow JL, et al. 2008. Variation analysis and gene annotation of eight MHC haplotypes: the MHC Haplotype Project. Immunogenetics 60: 1–18.

Howe K, Wood JM. 2015. Using optical mapping data for the improvement of vertebrate genome assemblies. Gigascience 4: 10.

Huddleston J, Ranade S, Malig M, Antonacci F, Chaisson M, Hon L, Sudmant PH, Graves TA, Alkan C, Dennis MY, et al. 2014. Reconstructing complex regions of genomes using long-read sequencing technology. Genome Res 24: 688–696.

International HapMap Consortium. 2005. A haplotype map of the human genome. Nature 437: 1299–1320.

International Human Genome Sequencing Consortium. 2004. Finishing the euchromatic sequence of the human genome. Nature 431: 931–945.

Kidd JM, Cooper GM, Donahue WF, Hayden HS, Sampas N, Graves T, Hansen N, Teague B, Alkan C, Antonacci F, et al. 2008. Mapping and sequencing of structural variation from eight human genomes. Nature 453: 56–64.

Kitts PA, Church DM, Thibaud-Nissen F, Choi J, Hem V, Sapojnikov V, Smith RG, Tatusova T, Xiang C, Zherikov A, et al. 2016. Assembly: a resource for assembled genomes at NCBI. Nucleic Acids Res 44: D73–80.

Lander ES, Linton LM, Birren B, Nusbaum C, Zody MC, Baldwin J, Devon K, Dewar K, Doyle M, FitzHugh W, et al. 2001. Initial sequencing and analysis of the human genome. Nature 409: 860–921.

Landrum MJ, Lee JM, Riley GR, Jang W, Rubinstein WS, Church DM, Maglott DR. 2014. ClinVar: public archive of relationships among sequence variation and human phenotype. Nucleic Acids Res 42: D980–5.

Levy S, Sutton G, Ng PC, Feuk L, Halpern AL, Walenz BP, Axelrod N, Huang J, Kirkness EF, Denisov G, et al. 2007. The diploid genome sequence of an individual human. PLoS Biol 5: e254.

Li H. 2013. Aligning sequence reads, clone sequences and assembly contigs with BWA-MEM. arXiv [q-bioGN]. http://arxiv.org/abs/1303.3997.

Li H. 2015. FermiKit: assembly-based variant calling for Illumina resequencing data. Bioinformatics 31: 3694–3696.

Li H. 2014. Toward better understanding of artifacts in variant calling from high-coverage samples. Bioinformatics 30: 2843–2851.

Li R, Li Y, Zheng H, Luo R, Zhu H, Li Q, Qian W, Ren Y, Tian G, Li J, et al. 2010. Building the sequence map of the human pan-genome. Nat Biotechnol 28: 57–63.

Matsuno H, Ohi K, Hashimoto R, Yamamori H, Yasuda Y, Fujimoto M, Yano-Umeda S, Saneyoshi T, Takeda M, Hayashi Y. 2015. A naturally occurring null variant of the NMDA type glutamate receptor NR3B subunit is a risk factor of schizophrenia. PLoS One 10: e0116319.

McKenna A, Hanna M, Banks E, Sivachenko A, Cibulskis K, Kernytsky A, Garimella K, Altshuler D, Gabriel S, Daly M, et al. 2010. The Genome Analysis Toolkit: a MapReduce framework for analyzing next-generation DNA sequencing data. Genome Res 20: 1297–1303.

Miga KH, Eisenhart C, Kent WJ. 2015. Utilizing mapping targets of sequences underrepresented in the reference assembly to reduce false positive alignments. Nucleic Acids Res 43: e133.

Miga KH, Newton Y, Jain M, Altemose N, Willard HF, Kent WJ. 2014. Centromere reference models for human chromosomes X and Y satellite arrays. Genome Res 24: 697–707.

Mueller JL, Skaletsky H, Brown LG, Zaghlul S, Rock S, Graves T, Auger K, Warren WC, Wilson RK, Page DC. 2013. Independent specialization of the human and mouse X chromosomes for the male germ line. Nat Genet 45: 1083–1087.

Nguyen N, Hickey G, Zerbino DR, Raney B, Earl D, Armstrong J, Kent WJ, Haussler D, Paten B. 2015. Building a pan-genome reference for a population. J Comput Biol 22: 387–401.

Novak AM, Garrison E, Paten B. 2016. A Graph Extension of the Positional Burrows-Wheeler Transform and its Applications. http://biorxiv.org/lookup/doi/10.1101/051409.

O’Leary NA, Wright MW, Brister JR, Ciufo S, Haddad D, McVeigh R, Rajput B, Robbertse B, Smith-White B, Ako-Adjei D, et al. 2016. Reference sequence (RefSeq) database at NCBI: current status, taxonomic expansion, and functional annotation. Nucleic Acids Res 44: D733–45.

Paten B, Novak A, Haussler D. 2014. Mapping to a Reference Genome Structure. arXiv [qbioGN]. http://arxiv.org/abs/1404.5010.

Pendleton M, Sebra R, Pang AWC, Ummat A, Franzen O, Rausch T, Stütz AM, Stedman W, Anantharaman T, Hastie A, et al. 2015. Assembly and diploid architecture of an individual human genome via single-molecule technologies. Nat Methods 12: 780–786.

Pierson E, GTEx Consortium, Koller D, Battle A, Mostafavi S, Ardlie KG, Getz G, Wright FA, Kellis M, Volpi S, et al. 2015. Sharing and Specificity of Co-expression Networks across 35 Human Tissues. PLoS Comput Biol 11: e1004220.

Pyo C-W, Guethlein LA, Vu Q, Wang R, Abi-Rached L, Norman PJ, Marsh SGE, Miller JS, Parham P, Geraghty DE. 2010. Different patterns of evolution in the centromeric and telomeric regions of group A and B haplotypes of the human killer cell Ig-like receptor locus. PLoS One 5: e15115.

Rosenfeld JA, Mason CE, Smith TM. 2012. Limitations of the human reference genome for personalized genomics. PLoS One 7: e40294.

Samocha KE, Robinson EB, Sanders SJ, Stevens C, Sabo A, McGrath LM, Kosmicki JA, Rehnström K, Mallick S, Kirby A, et al. 2014. A framework for the interpretation of de novo mutation in human disease. Nat Genet 46: 944–950.

Schmutz J, Martin J, Terry A, Couronne O, Grimwood J, Lowry S, Gordon LA, Scott D, Xie G, Huang W, et al. 2004. The DNA sequence and comparative analysis of human chromosome 5. Nature 431: 268–274.

Schneider VA, Chen HC, Clausen C, Meric PA, Zhou Z, Bouk N, Husain N, Maglott DR, Church DM. 2013. Clone DB: an integrated NCBI resource for clone-associated data. Nucleic Acids Res 41: D1070–8.

Sharp AJ, Locke DP, McGrath SD, Cheng Z, Bailey JA, Vallente RU, Pertz LM, Clark RA, Schwartz S, Segraves R, et al. 2005. Segmental duplications and copy-number variation in the human genome. Am J Hum Genet 77: 78–88.

Shi L, Guo Y, Dong C, Huddleston J, Yang H, Han X, Fu A, Li Q, Li N, Gong S, et al. 2016. Long-read sequencing and de novo assembly of a Chinese genome. Nat Commun 7: 12065.

Steinberg KM, Schneider VA, Graves-Lindsay TA, Fulton RS, Agarwala R, Huddleston J, Shiryev SA, Morgulis A, Surti U, Warren WC, et al. 2014. Single haplotype assembly of the human genome from a hydatidiform mole. Genome Res 24: 2066–2076.

Sudmant PH, Kitzman JO, Antonacci F, Alkan C, Malig M, Tsalenko A, Sampas N, Bruhn L, Shendure J, 1000 Genomes Project, et al. 2010. Diversity of human copy number variation and multicopy genes. Science 330: 641–646.

Teague B, Waterman MS, Goldstein S, Potamousis K, Zhou S, Reslewic S, Sarkar D, Valouev A, Churas C, Kidd JM, et al. 2010. High-resolution human genome structure by single-molecule analysis. Proc Natl Acad Sci U S A 107: 10848–10853.

Vezzi F, Narzisi G, Mishra B. 2012. Reevaluating assembly evaluations with feature response curves: GAGE and assemblathons. PLoS One 7: e52210.

Walker BJ, Abeel T, Shea T, Priest M, Abouelliel A, Sakthikumar S, Cuomo CA, Zeng Q, Wortman J, Young SK, et al. 2014. Pilon: an integrated tool for comprehensive microbial variant detection and genome assembly improvement. PLoS One 9: e112963.

Wang J, Wang W, Li R, Li Y, Tian G, Goodman L, Fan W, Zhang J, Li J, Zhang J, et al. 2008. The diploid genome sequence of an Asian individual. Nature 456: 60–65.

Watson CT, Steinberg KM, Graves TA, Warren RL, Malig M, Schein J, Wilson RK, Holt RA, Eichler EE, Breden F. 2015. Sequencing of the human IG light chain loci from a hydatidiform mole BAC library reveals locus-specific signatures of genetic diversity. Genes Immun 16: 24–34.

Watson CT, Steinberg KM, Huddleston J, Warren RL, Malig M, Schein J, Willsey AJ, Joy JB, Scott JK, Graves TA, et al. 2013. Complete haplotype sequence of the human immunoglobulin heavy-chain variable, diversity, and joining genes and characterization of allelic and copy-number variation. Am J Hum Genet 92: 530–546.

Xue Y, Sun D, Daly A, Yang F, Zhou X, Zhao M, Huang N, Zerjal T, Lee C, Carter NP, et al. 2008. Adaptive evolution of UGT2B17 copy-number variation. Am J Hum Genet 83: 337–346.

Zhao H, Sun Z, Wang J, Huang H, Kocher J-P, Wang L. 2014. CrossMap: a versatile tool for coordinate conversion between genome assemblies. Bioinformatics 30: 1006–1007.

Zody MC, Jiang Z, Fung H-C, Antonacci F, Hillier LW, Cardone MF, Graves TA, Kidd JM, Cheng Z, Abouelleil A, et al. 2008. Evolutionary toggling of the MAPT 17q21. 31 inversion region. Nat Genet 40: 1076–1083.

Zook JM, Catoe D, McDaniel J, Vang L, Spies N, Sidow A, Weng Z, Liu Y, Mason CE, Alexander N, et al. 2016. Extensive sequencing of seven human genomes to characterize benchmark reference materials. Sci Data 3: 160025.

Zook JM, Chapman B, Wang J, Mittelman D, Hofmann O, Hide W, Salit M. 2014. Integrating human sequence data sets provides a resource of benchmark SNP and indel genotype calls. Nat Biotechnol 32: 246–251.

